# Evaluating High Spatial Resolution Diffusion Kurtosis Imaging at 3T: Reproducibility and Quality of Fit

**DOI:** 10.1101/2020.07.10.197921

**Authors:** Loxlan W. Kasa, Roy A.M. Haast, Tristan K. Kuehn, Farah N. Mushtaha, Corey A. Baron, Terry Peters, Ali R. Khan

**Affiliations:** Imaging Research Laboratories, Robarts Research Institute, London, Ontario, Canada; School of Biomedical Engineering, Western University, London, Ontario, Canada; Centre for Functional and Metabolic Mapping, Robarts Research Institute, Western University, London, Ontario, Canada; Department of Medical Biophysics, Western University, London, Ontario, Canada; Departments of Medical Imaging, Western University, London, Ontario, Canada

**Keywords:** Diffusion magnetic resonance imaging, diffusion kurtosis imaging, reproducibility, quality of fitting

## Abstract

**Background:** Diffusion kurtosis imaging (DKI) quantifies the microstructure’s non-Gaussian diffusion properties. However, it has increased fitting parameters and requires higher b-values. Evaluation of DKI reproducibility is important for clinical purposes.

**Purpose:** To assess reproducibility in whole-brain high resolution DKI at varying b-values.

**Study Type:** Prospective.

**Subjects and Phantoms:** Forty-four individuals from the test-retest Human Connectome Project (HCP) database and twelve 3D-printed tissue mimicking phantoms.

**Field Strength/Sequence:** Multiband echo-planar imaging for *in vivo* and phantom diffusion-weighted imaging at 3T and 9.4T respectively. MPRAGE at 3T for *in vivo* structural data.

**Assessment:** From HCP data with b-value =1000,2000,3000 s/mm^2^ (dataset A), two additional datasets with b-values=1000, 3000 s/mm^2^ (dataset B) and b-values=1000, 2000 s/mm^2^ (dataset C) were extracted. Estimated DKI metrics from each dataset were used for evaluating reproducibility and fitting quality in whole-brain white matter (WM), region of interest (ROI) and gray matter (GM).

**Statistical Tests:** DKI reproducibility was assessed using the within-subject coefficient of variation (CoV), fitting residuals to evaluate DKI fitting accuracy and Pearson’s correlation to investigate presence of systematic biases.

**Results:** Compared to dataset C, the CoV from DKI parameters from datasets A and B were comparable, with WM and GM CoVs <20%, while differences between datasets were smaller for the DKI-derived DTI parameters. Slightly higher fitting residuals were observed in dataset C compared to A and B, but lower residuals in dataset B were detected for the WM ROIs. A similar trend was observed for the phantom data with comparable CoVs at varying fiber orientations for datasets A and B. In addition, dataset C was characterized by higher residuals across the different fiber crossings.

**Data Conclusion:** The comparable reproducibility of DKI maps between datasets A and B observed in the *in vivo* and phantom data indicates that high reproducibility can still be achieved within a reasonable scan time, supporting DKI for clinical purposes.

**HIGHLIGHTS:**

I. Reproducibility and fitting accuracy of high resolution DKI were evaluated as function of available b-values.
II. A DKI dataset with b-values of 1000 and 3000 s/mm^2^ performs equally well as the original HCP three-shell dataset, while a dataset with b-values of 1000 and 2000 s/mm^2^ has lower reproducibility and fitting quality.
III. *In vivo* results were verified using phantoms capable of mimicking different white matter configurations.
IV. These results suggest that DKI data can be obtained within less time, without sacrificing data quality.

## 1. INTRODUCTION

Diffusion tensor imaging (DTI) is an elegant noninvasive method to estimate water particle’s apparent diffusion in the brain (1). Since its first use (1), DTI has evolved tremendously, including faster acquisition time, making it better suitable for diagnostic purposes (2). Common quantitative parameters derived from DTI include: fractional anisotropy (FA), which describes the amount of anisotropic diffusion and reflects the preferred direction of diffusion; mean diffusivity (MD), the average diffusion; axial diffusivity (AD), the diffusion in the axial direction; and radial diffusivity (RD), the diffusion perpendicular to the axial diffusion (3). While these DTI metrics are useful, the technique is based on the assumption that diffusion of water particles in neuronal tissue follows a Gaussian distribution. Although this is more true in regions of coherent fiber bundles, DTI fails to adequately quantify most parts of the brain with complex cytoarchitecture such as in the cerebral cortex and in areas of white-matter (WM) with substantial fiber crossings (3). These microstructural properties cause water diffusion to considerably deviate from a Gaussian shape. This has been evident in imaging techniques like *q*-space imaging, which employs many acquisitions that include both high and low b-values in order to estimate the full diffusion displacement but with the trade-off of longer scan time (3, 4).

Diffusion kurtosis imaging (DKI) was introduced to accommodate the shortcomings of DTI and better characterize non-Gaussian diffusion behaviour with a protocol that uses a modest increase in b-value over that used in DTI (5). DKI is derived from expanding the standard diffusion signal equation in powers of b-value (see Eq.1). Therefore, both diffusion and kurtosis tensors can be estimated (5, 6) to provide mean kurtosis (MK); axial kurtosis (AK); radial kurtosis (RK) in addition to the DTI metrics. Since the introduction of these parameters, they have been used across several clinical populations including Parkinson’s and Alzheimer’s patients (7), but also for (early) assessment of stroke (8), traumatic brain injury (9), epilepsy (10) and numerous other clinical studies.

Despite the benefits of DKI as a non-invasive tool for delineating microstructural alterations due to diseases or microstructural complexity, the technique requires at least one additional shell of acquisitions than the single shell required for DTI. DKI also has more parameters to fit than DTI, which, depending on the acquisition (e.g., minimum of 15 directions for standard DKI protocol), could lead to overfitting of the data, poor reproducibility (test-retest reliability), and limited use in clinical practice. Additionally, in order to capture the non-Gaussian diffusion behaviour of water molecules in biological tissues, b-values larger than those employed in DTI are required (5). However, higher b-values not only lower the signal-to-noise ratio (SNR) of the respective image volumes, affecting the reproducibility of the calculated parameters, but also increase acquisition time (11–13). In order for DKI to be integrated into clinical workflows, the reproducibility of its estimated parameters in different tissue types with different microstructural properties has to be established.

A previous study has assessed DKI reproducibility with different b-values and fitting algorithms, however was limited to 3 mm isotropic spatial resolution, and concentrated on MK in selected WM and gray matter (GM) voxels only (14). A more recent study specifically focussed on test-retest reliability of high spatial resolution data, looked at test-retest reliability of DKI 1.75 mm isotropic spatial resolution only, but were unable to utilize 1.25 mm resolution imaging due to insufficient SNR (15). Although this was a first study to evaluate DKI at high spatial resolution *in-vivo*, the study only focused on whole brain WM and the effect of different b-values in different tissue types was not assessed. Since DKI has the ability to characterize microstructure in less anisotropic environments (5) such as GM, it is important to fully investigate DKI reproducibility in these areas which could also aid our understanding of neurodegenerative diseases including aging (16, 17) that are associated with GM abnormalities. The aim of the study is therefore to evaluate the test-retest reliability of high resolution (1.25 mm isotropic spatial resolution) DKI estimated parameters over the entire brain, and specifically investigate whether acquisitions with fewer shells, but differing maximum b-value, could provide reproducible parameters with reduced scanning time. In addition, we expect this relation to vary across structures based on the expected non-Gaussian diffusion profiles induced by differences in their microstructural composition. Therefore, we explicitly analyse test-retest reliability of cortical GM, whole-brain WM and in selected WM regions of interest (ROIs), i.e., fiber bundles known to be implicated in neurological disorders (18). Moreover, we verify the reproducibility of each DKI parameter with fibre phantoms that can mimic common WM fiber configurations, including variable degrees of crossing fibers (19).

## 2. MATERIALS AND METHODS

### 2.1 Image acquisition

#### In vivo Imaging

A total of 44 subjects (31 female, age 22 to 35) from the test-retest Human Connectome Project (HCP) database were included in this study. Diffusion weighted imaging (DWI) data were acquired twice for all subjects across two separate sessions (average of 5 ± 3 months apart) using a high-quality image acquisition protocol and a modified Siemens Skyra 3T scanner (20). DWI acquisition parameters included TR/TE=5520/89.5 ms, multiband factor = 3, phase Partial Fourier = 6/8 without in-plane acceleration, and nominal isotropic voxel size of 1.25 mm^3^. A total of 288 images were acquired in each DWI dataset (acquired in both left-to-right and right-to-left phase-encoding polarities for EPI distortion correction), including 18 baseline images with a low diffusion weighting b-value =5 s/mm^2^ and 270 diffusion weighted images evenly distributed across three shells of b-values =1000, 2000, 3000 s/mm^2^. We made use of data pre-processed following HCP’s ‘minimal preprocessing’ pipeline (21), which included brain masking, motion correction, eddy current correction and EPI distortion correction.

#### Phantom Imaging

We constructed four groups of three phantoms each to compare across specific crossing angles of 0°, 30°, 60°, and 90°, respectively, and to verify the *in vivo* results. For this we used a 3D printing protocol developed in-house (19) that entails fused deposition modeling 3D printing with a composite material consisting of rubber-elastomeric polymer and a polyvinyl alcohol (PVA) component (PORO-LAY). Each of the 12 phantoms were 11 mm in radius with 100 μm layers of parallel lines with alternating orientations mimicking brain microstructure with crossing fibres. Following immersion in water for 168 hours to allow the PVA to dissolve exposing the microstructure, the phantoms were stacked in a test tube with distilled water before imaging with a 9.4T Bruker scanner, following the HCP *in vivo* protocol’s number of directions per shell. The other imaging parameters include TE/TR=37/2500 ms, FOV=200×200 mm^2^, 0.7 mm isotropic in-plane resolution, 6 axial slices (3 mm, one per phantom), and scan of time 8.5 min per each of 2 scans to cover all phantoms. This was repeated twice to facilitate test-retest reliability measurements (the sample was not removed between scans).

### 2.2 Image Processing

#### In vivo

For both test and retest data of each subject, two subsets of data were selected from the original three shell dataset to assess DKI reproducibility as a function of b-values used. The second dataset included only b-values = 1000, 3000 s/mm^2^, while a third only b-values = 1000, 2000 s/mm^2^. Note that the b-value = 0 s/mm^2^ was included in both datasets. Three separate fitting procedures were conducted using the open source software diffusion imaging in python (DIPY v1.0) (22) to generate corresponding DKI parametric maps for each b-value dataset and time point (i.e., test-retest). It is important to note that during DKI fitting, the diffusion tensor *D* and diffusion kurtosis *K* are calculated simultaneously (see Eq.1), where *K* characterizes the deviation from Gaussian diffusion (5). *S* corresponds to the diffusion weighted image at *b* ≠ 0 and *S*_0_ is non-diffusion weighted image at *b* = 0.

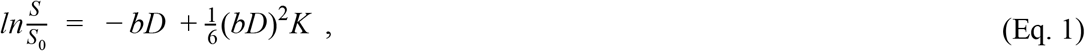

The DKI metrics include: the DTI MD, AD and RD. In addition, we obtained the three kurtosis metrics: MK, the average of the kurtosis over all diffusion directions; AK, the kurtosis in the axial direction and RK, the kurtosis perpendicular to the axial direction. To allow group-analyses, we used the MRtrix v3.0 (23) functions, ‘*dwi2fod*’ to generate individual fiber orientation dispersion (FOD) maps and ‘*population_template*’ to compute an unbiased group-average FOD template from the individual subjects’s FODs. To minimize variability between the subjects’ scans (test and retest data), we performed a rigid registration between subject’s FODs bringing all FODs into test data space. The calculated transformations were also applied to the respective DKI maps. Then the FODs from the first test dataset for each subject were registered to the FOD template, the calculated warps were then used to warp each subject’s DKI maps including maps from retest data in test data space to the template space. For white matter region-based analyses, the JHU-ICBM-labels atlas (24) in MNI152 space was transformed to the FOD template via transformation obtained from FSL v6.0.2 ‘*flirt*’ registration of the MNI (6^th^ generation) template to b=0 image extracted from the FOD template. The MNI152 to FOD template transformation was also used for transforming selected white-matter ROIs from the JHU-ICBM-labels atlas. We selected these white-matter ROIs (see supplementary data Figure S1), (CC-Cingulum Cingulate, CH-Cingulum Hippocampus, Fx-Fornix, SFOF-Superior Fronto Occipital Fasciculus, SLF-Superior Longitudinal Fasciculus and Unc-Uncinate Fasciculus) since they are commonly implicated in neurological disorders (18). In addition, for grey matter (GM) analyses of the individual lobes (FL-Frontal Lobe, PL-Parietal Lobe, LL-Limbic Lobe, TL-Temporal Lobe and OL-Occipital Lobe) of the population-average, landmark and surface-based (PALS) atlas of human cerebral cortex (25), we mapped all the individual subject’s co-registered maps onto the ‘fsaverage’ surface space (26). To do so we used the DWI-based data that were registered to the anatomical space following the HCP’s ‘minimal preprocessing’ pipeline (21). Then, FreeSurfer’s ‘*mri_vol2surf*’ function was used to project the anatomical space DKI maps onto the fsaverage surface by calculating the vertex-wise (i.e., per individual data point on the surface mesh) averages across cortical depths (i.e., WM to pial surface direction). To minimize partial voluming effects with WM and CSF, we only averaged across voxels within 20 to 80 percent of the estimated cortical thickness, within each of the five lobes from the PALS atlas (see supplementary data Figure S1).

#### Phantoms

For the phantom’s test-retest data, we manually created masks for the images from each of the 12 phantoms to remove regions with air bubbles. Then the original test-retest data were used for selecting two subsets of data with two shells each, similar to the procedure performed for the *in-vivo* dataset. Following this, FODs were generated for each subset including the original data with three shells. We then rigidly registered the retest to the test data to minimize any variability between the scans, and the warp fields obtained were used to transform the calculated DKI maps. These steps were also similar to the *in-vivo* dataset workflow, with the exception of the transformation to the WM and GM template spaces.

### 2.3 Image Analysis

To evaluate reproducibility, we calculated the voxel- (both in vivo and phantom data) and vertex-wise (in vivo data only) within-subject coefficient of variation (CoV). The CoV (i.e., the standard deviation to mean ratio) was determined for the individual parametric maps and estimated from the test-retest data within each of the three datasets: original (b-value = 1000, 2000, 3000 s/mm^2^) and the two selected datasets (b-value = 1000, 3000 s/mm^2^ and b-value = 1000, 2000 s/mm^2^). In addition, to check for any potential bias in our analysis, we performed Pearson correlation tests of the calculated values between the DKI parametric maps derived from each datasets (27). Moreover, to evaluate DKI fitting quality, we calculated the mean absolute residuals (*R*) (3), which represent the difference between the modelled and diffusion weighted signals (see Eq. 2) for each dataset (i.e., varying b-values). Large residuals indicate the presence either of potential artefacts in the acquired data, or that the applied method is unable to characterize the observed signal accurately (3). Therefore, to gauge the quality of the DKI estimation of the measured signals, mean residual maps *R* for each voxel for individual DWIs were calculated (28),

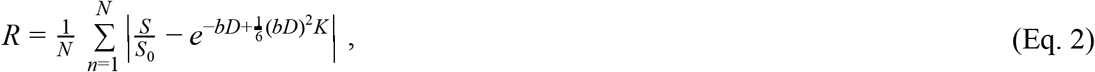

where *N* is the number DWIs, *S* and *S*_0_ are the diffusion and non-diffusion weighted images respectively, and *D* and *K* are the estimated diffusion and kurtosis tensor with the diffusion weighting *b*. Note that in addition to *K*, *D* is also estimated in the kurtosis method, as outlined by Jensen et al. (5). All statistical analysis was carried out using the scipy package (v1.4.1)for Python.

**Figure 1.**
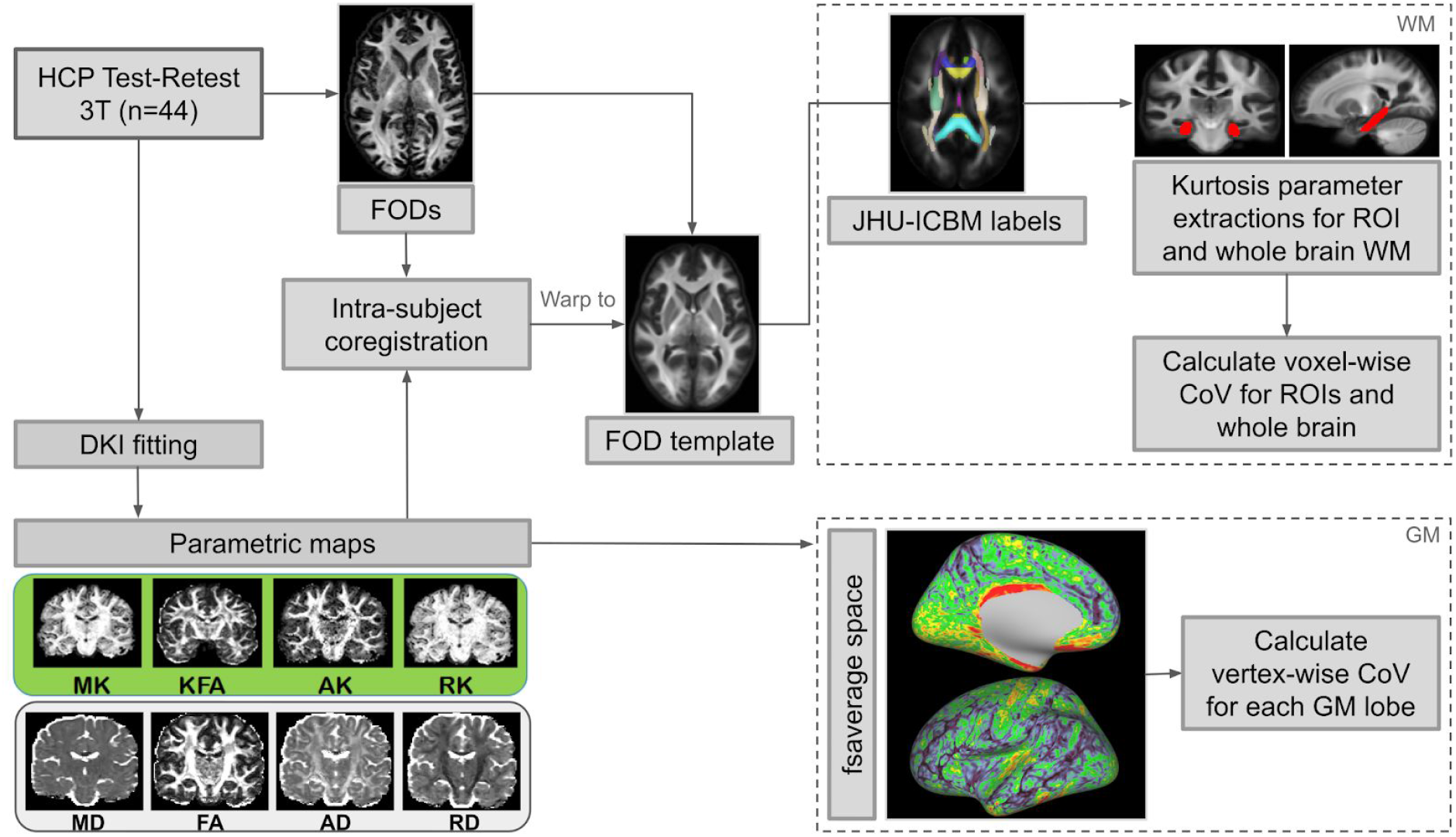
General workflow. Preprocessed HCP test-retest data were used for the study. A population-based template was created from the subjects’ FOD images. Then, the individual subject’s generated DKI maps were co-registered before warping into the FOD template space. The WM JHU-ICBM-labels atlas was co-registered to the template, facilitating CoV analysis in WM ROIs and whole-brain WM. For GM analysis, the co-registered subject’s maps are mapped onto Freesurfer’s fsaverage space, following this CoV analysis was performed for selected ROI’s surfaces.

**Figure 2.**
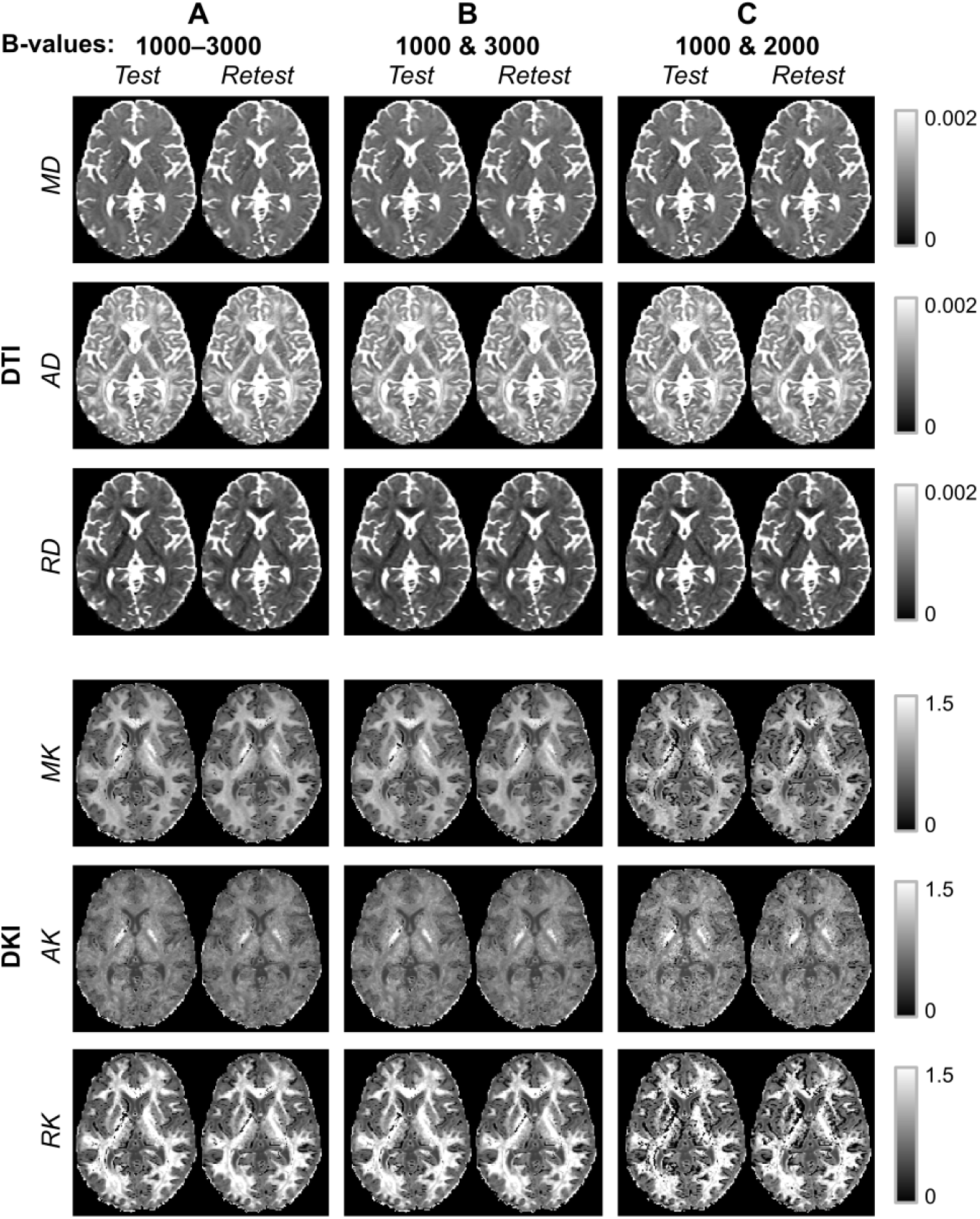
The test-retest datasets, (b-values = 1000-3000 s/mm^2^) 1^st^ column (dataset A) and the two derived datasets (b-values = 1000 & 3000) 2^nd^ column (dataset B) and (b-values = 1000 & 2000 s/mm^2^) 3^rd^ column (dataset C). Showing is a representative axial slice for all generated DKI maps including the DTI maps estimated with DKI method from an individual subject.

## 3. RESULTS

The rest of the section is divided into two subsections: *in vivo* imaging and phantom results. For the *in-vivo* study, Figures 3 and 4 report the CoV analysis of the WM while Figure 5 and 6 represent GM CoV analysis and evaluation of whole-brain and WM ROIs fitting quality, respectively. Figures 7 and 8 gives us the CoV and fitting quality analysis of the phantom data.

**Figure 3.**
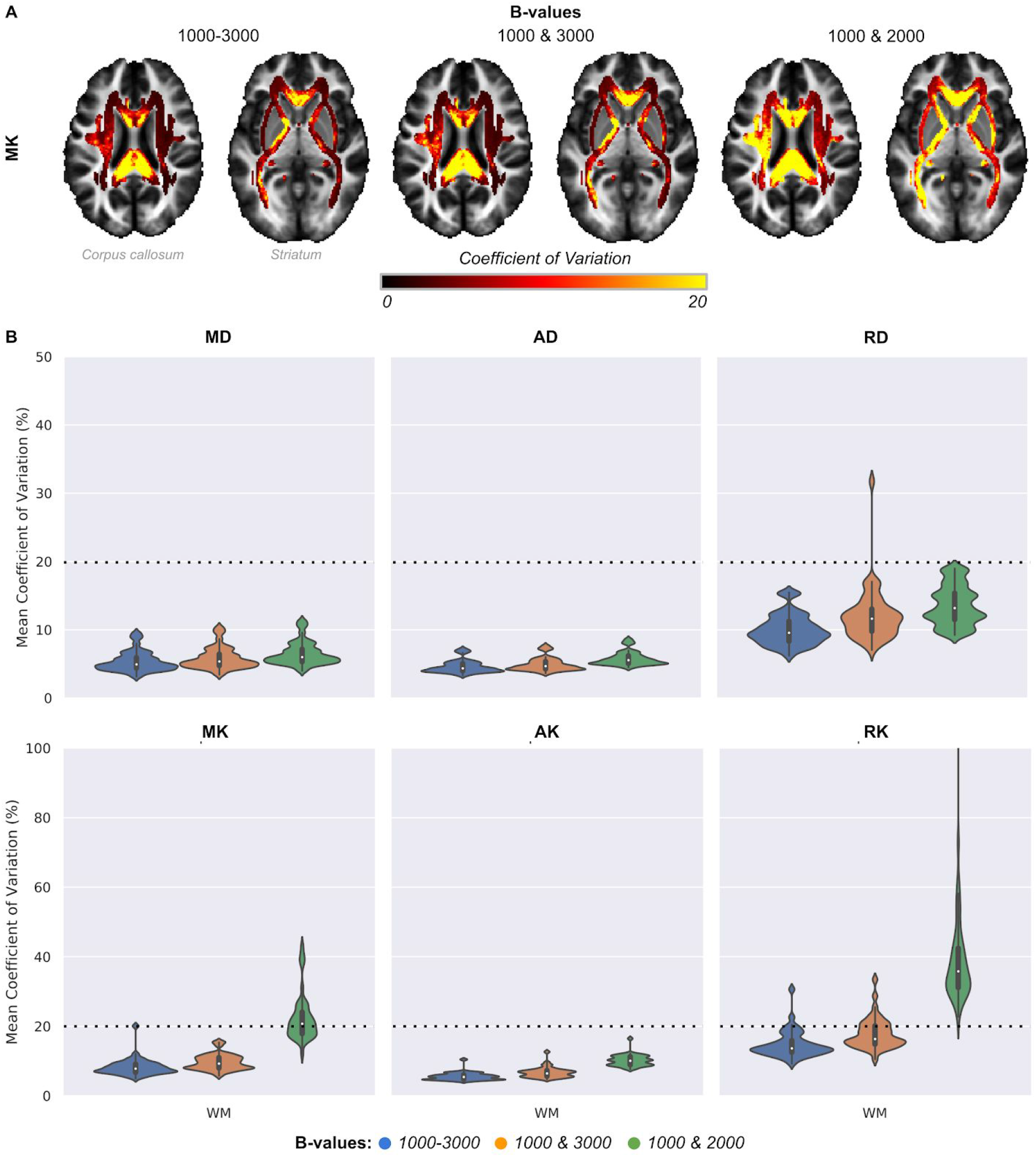
Mean voxel-wise within-subject CoV for MK mapped onto the FOD template (A) and within the WM for all the maps (B). All maps were generated from dataset A (blue) (b-value = 1000-3000 s/mm^2^), dataset B(orange) (b-value = 1000 & 3000 s/mm^2^) and dataset C (green) (b-value = 1000 & 2000 s/mm^2^). The dotted line is placed at 20% CoV for reference.

**Figure 4.**
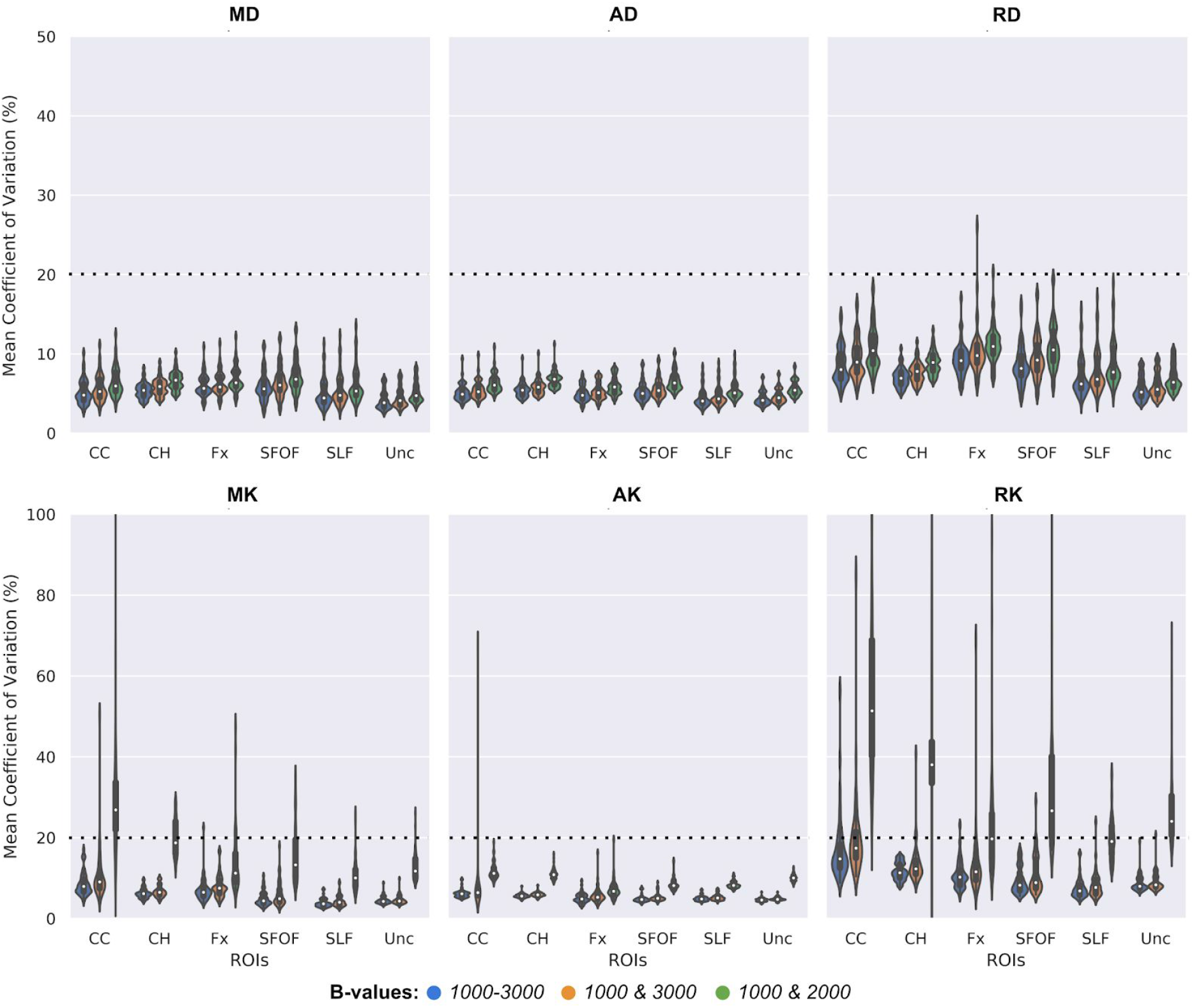
Mean voxel-wise within-subject CoV within the WM ROIs (CC-Cingulum Cingulate, CH-Cingulum Hippocampus, Fx-Fornix, SFOF-Superior Fronto Occipital Fasciculus, SLF-Superior Longitudinal Fasciculus and Unc-Uncinate Fasciculus). All maps were generated from the datasets; dataset A (blue) (b-value = 1000-3000 s/mm^2^), dataset B (orange) (b-value = 1000 & 3000 s/mm^2^) and dataset C (green) (b-value = 1000 & 2000 s/mm^2^). The dotted line is placed at 20% CoV for reference.

**Figure 5.**
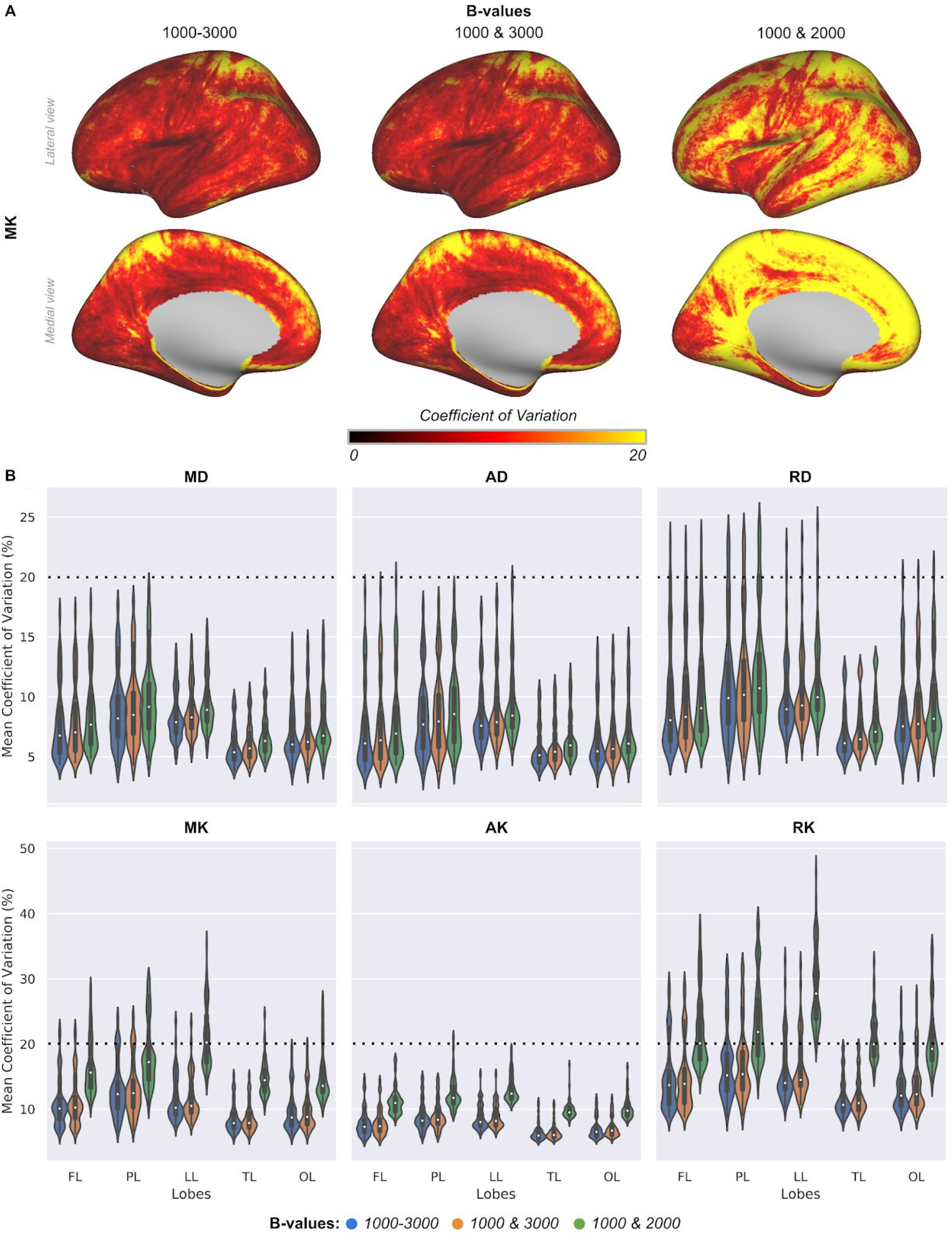
Mean vertex-wise within-subject coefficient of variation: for MK mapped onto fsaverage (A) and all DKI maps within the five lobes (FL-Frontal Lobe, PL-Parietal Lobe, LL-Limbic Lobe, TL-Temporal Lobe and OL-Occipital Lobe) (B). The maps were generated from the datasets; dataset A (blue) (b-value = 1000-3000 s/mm^2^), dataset B (orange) (b-value = 1000 & 3000 s/mm^2^) and dataset C (green) (b-value = 1000 & 2000 s/mm^2^). Showing for the left hemisphere only since a similar trend was observed in the right hemisphere.The dotted line is placed at 20% CoV for reference.

**Figure 6.**
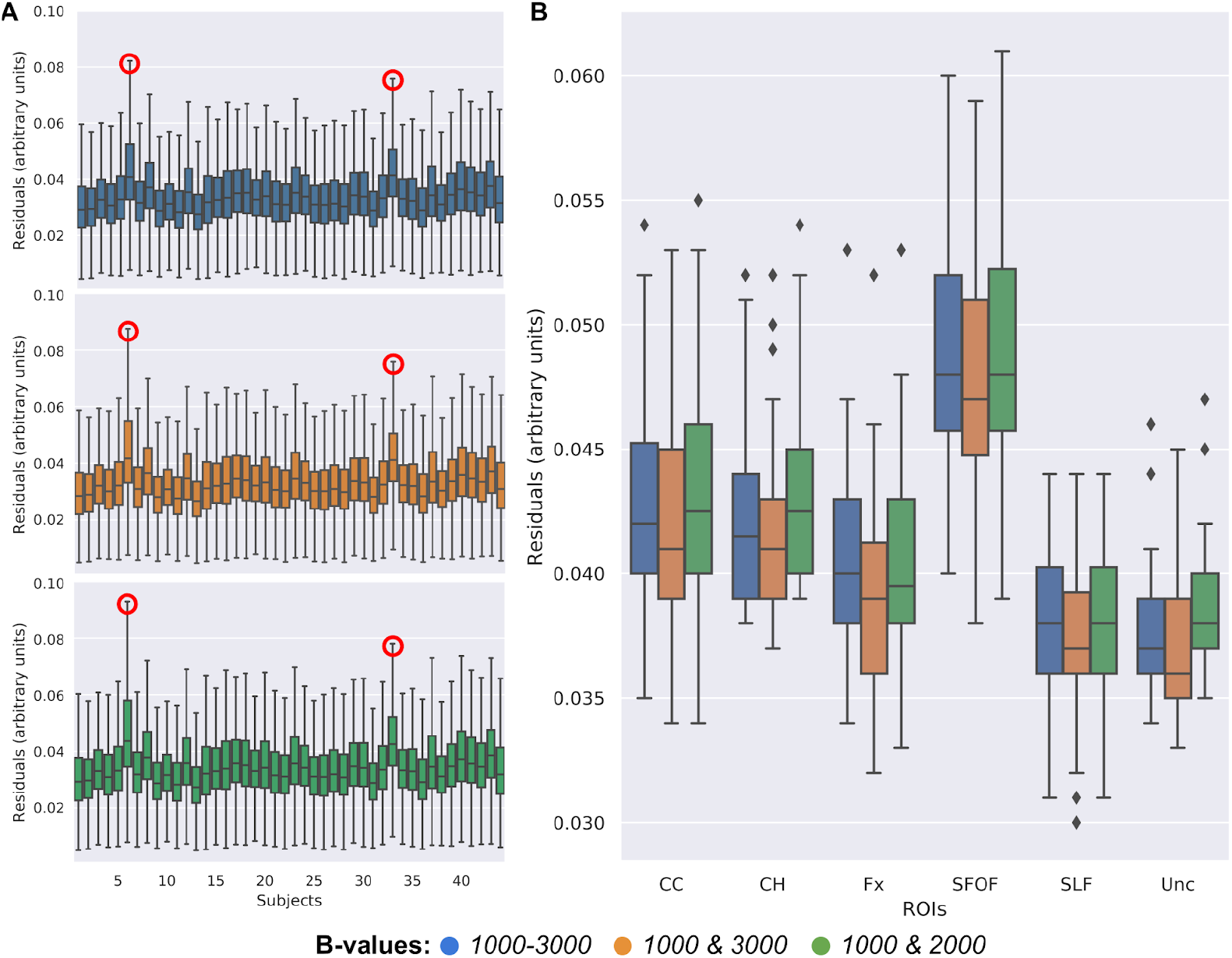
Residuals from DKI fitting averaged across voxels for individual subject’s DWIs from each of the three datasets:A (blue) (b-value = 1000-3000 s/mm^2^), dataset B (orange) (b-value = 1000 & 3000 s/mm^2^) and dataset C (green) (b-value = 1000 & 2000 s/mm^2^) (A). ROI residuals from DKI fitting averaged across voxels for individual DWIs from each of the three datasets: A, B and C (B).

**Figure 7.**
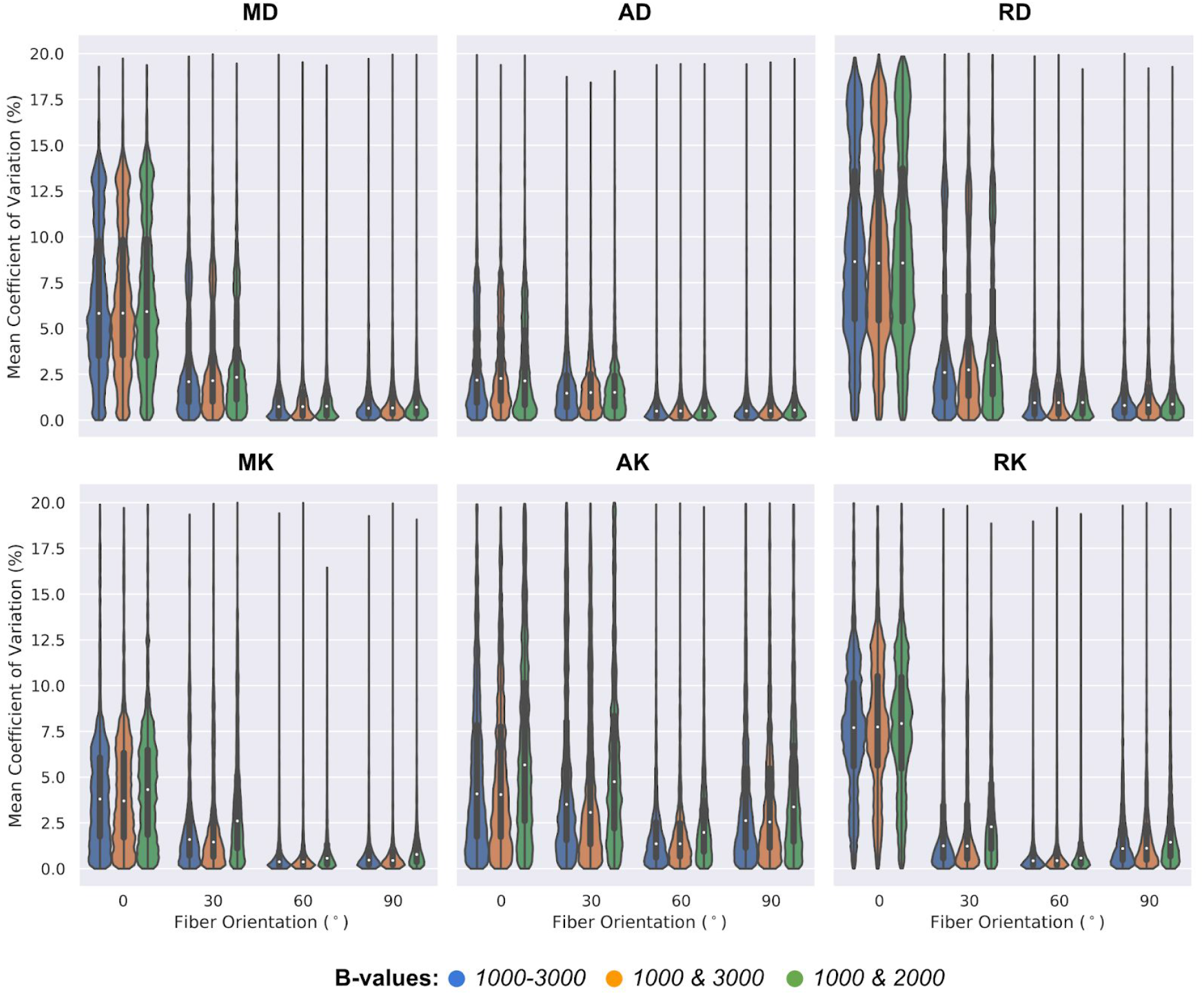
Mean voxel-wise within-scans coefficient of variation determined across the phantoms with varying fiber crossings (0°, 30°, 60° and 90°). All maps generated from the dataset A (blue) (b-value = 1000-3000 s/mm^2^) and the derived datasets, B (orange) (b-value = 1000 & 3000 s/mm^2^) and C (green) (b-value = 1000 & 2000 s/mm^2^) respectively.

**Figure 8.**
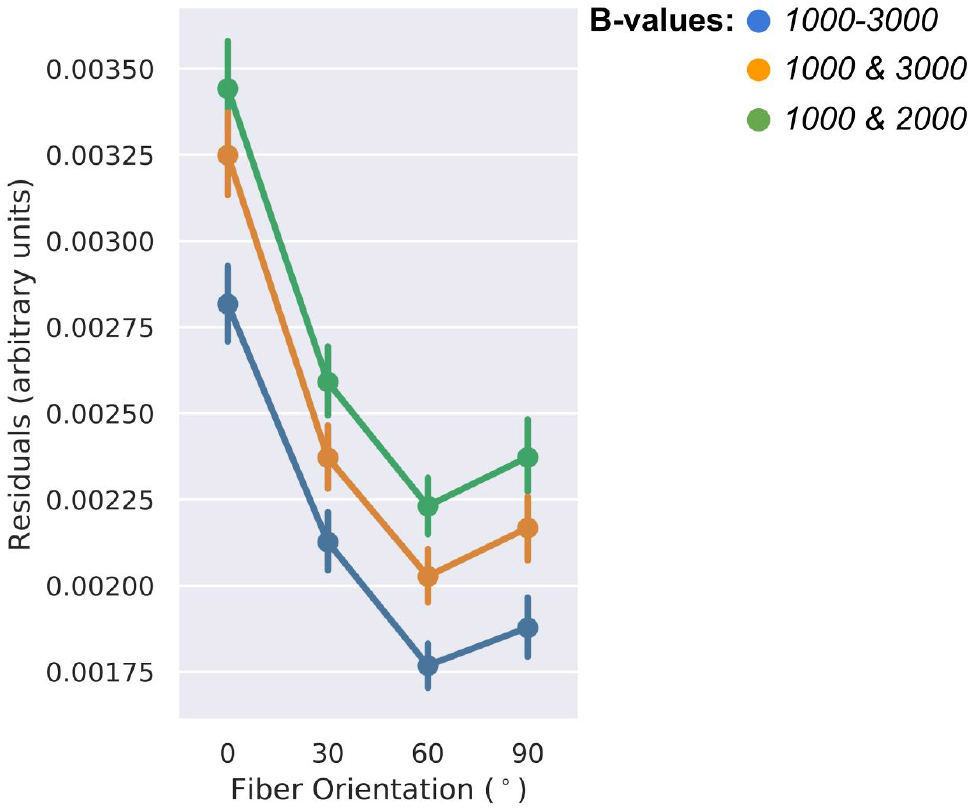
Fitting residuals averaged across voxels for individual DWIs in each of the phantoms with varying fiber crossings (0°, 30°, 60° and 90°). This is determined for each of the three datasets, A (blue) (b-value = 1000-3000 s/mm^2^), B (orange) (b-value = 1000 & 3000 s/mm^2^) and C (geen) (b-value = 1000 & 2000 s/mm^2^) respectively.

### 3.1. In vivo Imaging

#### Parameter reproducibility

Figure 2 shows a representative axial slice for all the DKI estimated parameters from the test-retest HCP three shells dataset (1^st^ column) from here on will be dataset A and the derived two shells datasets, will be referred to as dataset B (2^nd^ column) and dataset C (3^rd^ column). From visual judgement, we notice a close resemblance in the DKI parametric maps (MK, AK and RK) between datasets A and B, but not dataset C. However, not much difference is seen in DTI maps calculated from the DKI technique.

The heat maps in Figure 3A indicate the variation in MK metric within the whole-brain WM mask. The CoV for datasets A and B is comparable, while that for C differs significantly from the other two, with more regions around the ventricles having higher CoV (~ 20%). Moreover, the test-retest variability between the datasets A (in blue) and B (orange) were more comparable with mean CoV (< 20%) compared to C (green) for all the DKI metrics (MK, AK and RK) as shown in Figure 3B, though less differences is seen among the DTI metrics (MD, AD and RD) estimated from DKI. A similar trend was also seen from datasets A and B within-subject CoV in the selected WM ROIs as shown in Figure 4. In both datasets, the mean CoV ranges from 5-15% compared to dataset C with a more dispersed CoV as high as ~40% seen in the RK metric. In addition, we note a high Pearson correlation coefficient (r~1) between all the three datasets in their DKI mean parametric values in the whole-brain and selected WM ROIs as shown in the supplementary data (Figure S2–S3).

The heat maps in Figure 5A indicate the within-subject variation for the MK cortical surface data and show a similar behaviour as for the WM results (Figure 3). We observed widespread high CoV (~20%) in the cortex from the dataset C, in contrast to the A and B. This pattern is consistent across each of the five lobes, (Figure 5B). The CoV between the datasets A and B are similar in contrast to the dataset C. In terms of inter-regional differences in the CoV, higher variability ranging from (10-28%) is observed in the DKI maps (MK, AK and RK) with the highest in the limbic lobe from the dataset C compared to the A and B datasets with CoV (5-15%). Contrary to the high CoV in dataset C in DKI maps, a comparable reproducibility is observed in the estimated DTI maps from all the three datasets. Although there were differences in the CoV values across the three datasets (A, B and C), we note high Pearson correlation coefficient (r~1) between all the three datasets in their DKI mean parametric values as shown in supplementary data Figure S4.

#### Quality of fitting

Figure 6A shows the kurtosis tensor residuals calculated for each subject’s DWIs (averaged across whole-brain voxels). Two subjects (encircled in red) have high residuals across the three datasets with dataset C characterized by slightly higher residuals. Figure 6B presents the residuals averaged within the selected WM ROIs. We observe lower residuals in dataset B compared to A and C. Looking at individual ROIs DKI fitting, the SFOF had the highest fitting residuals compared to all the ROIs.

### 3.2 Phantom Studies

#### Parameter reproducibility

In line with the *in vivo* CoV analyses (Figures 3 and 4), CoV is lower for the datasets A and B compared to the C dataset for all DKI maps while DTI maps were almost identical in their mean CoV values from all three datasets, as shown in Figure 7. Interestingly, although dataset C values were consistently higher across the DKI maps across the different phantoms (i.e., characterized by different crossing angles), the lowest CoV was observed for the 60° phantom in all the maps from the respective datasets. The mean parametric values for MK and MD are shown in supplementary data Table S1, we observed all datasets having high Pearson correlation coefficient (r~1) except for 60° phantom in dataset C with an outlier.

#### Quality of fitting

The goodness of fit test for each of the phantoms representing varying fiber orientations is shown in Figure 8, where we observe the same trend with datasets A and B having lower fitting residuals compared to dataset C. Finally, the phantoms with 60° crossing angles had the lowest DKI fitting residuals across all the three datasets, similar to what is seen in Figure 7.

## 4. DISCUSSION

DKI has shown the ability to better characterize the brain’s microstructural composition by expanding the diffusion signal to estimate the kurtosis of the displacement distribution, in addition to the standard DTI-derived parameters (5, 29). This technique is seeing an increasing range of applications (30). However, DKI typically requires the addition of at least one additional b-value than required for traditional DTI. The higher b-value data have lower SNR, potentially affecting reproducibility, as well as increasing acquisition time, limiting clinical applicability. Therefore, we explored the reproducibility and fitting quality for all DKI-based estimated parameters at varying b-values used in a test-retest scenario for high spatial resolution data acquisitions. The key finding in this study was that DKI using only two b-values – the lowest and highest b-values (1000, 3000 s/mm^2^) of the HCP dataset – is a satisfactory approach. In other words, an additional third b-value (i.e., b-value = 2000 s/mm^2^) has a limited beneficial effect on the reproducibility to quantify the different tissue types (WM and GM). These findings were further verified with 3D printed brain microstructure-mimicking phantoms. Importantly, compared to the b-value = 1000, 2000, 3000 s/mm^2^ approach, the derived b-value = 1000, 3000 s/mm^2^ protocol offers the advantage of reduced scanning with comparable reproducibility.

### 4.1 DKI reproducibility and fitting quality in *in vivo* imaging

In the WM, the CoV for the DKI estimated parameters (MK, AK, RK, MD, AD and RD) was comparable between the original dataset with three shells and the third dataset with two shells (b-value=1000, 3000 s/mm^2^), with CoVs < 20%. This finding corresponds to a previous study which looked only at MK reproducibility and accuracy in selected voxels with respect to b-values and fitting algorithms in *in vivo* and *ex vivo* datasets (14). The authors concluded that in the WM, the protocols with maximum b-values (b_*max*_) ranging from 2500 to 3000 s/mm^2^ and applying weighted linear least square (WLS) fitting achieved the highest accuracy. It was also observed that in selected voxels, the variability of the microstructure influenced the accuracy and reproducibility for each b_*max*_. This is in line with what we examined in both whole-brain WM and across selected WM bundles. We also observed that there was lower variability within most of the selected bundles, including the larger known bundles like SLF and Unc for the datasets with b_*max*_ = 3000 s/mm^2^ (datasets A and B) compared to b_*max*_ = 2000 s/mm^2^ dataset (dataset C), except for a slight increase of CoV in the CC. These larger bundles are known to have higher crossing or bending configurations (3), suggesting that higher b-value acquisition could be beneficial for DKI to quantify non-Gaussian diffusion in WM regions with more complex fiber configurations. However, the SFOF results deviate from this hypothesis, with higher fitting residuals (i.e., lower quality of DKI fitting). This could be reflective of the label’s specific location, since SFOF is more coherent in the middle segment and disperses at the ends (31). Moreover, better representation of SFOF is seen to be achieved using data-driven approaches like tractography (31), therefore, the high residuals could be related to the inaccuracy of hand-segmented labels. For the GM, we focused on the five lobes and determined the average DKI coefficient of variation across cortical depths (i.e., from WM to the pial surface). Although DKI reproducibility was lower in GM compared to whole-brain WM, potentially due to partial volume effects (32, 33), a similar trend was noticed between the dataset A and the dataset B, both achieving CoVs ranging from 5-15%. Dataset C had higher inter-scan variability across the five lobes in all the maps. In contrast to the frontal, parietal, occipital and limbic lobes, the temporal lobe has lower CoV across all the DKI maps. This might allude to the structural composition of the temporal lobe itself compared to the other four lobes of the brain (25). In addition, possible signal drop outs in this may have could introduce consistently low values contributing to less variation between test-retest data. Furthermore, the differences in the reproducibility of each datasets across the lobes, in particular between dataset C and the two b_*max*_ = 3000 s/mm^2^ datasets (A and B), could be also due to the variability of the cytoarchitecture in individual lobes composition and functions (34).

To verify the quality of kurtosis tensor fitting, we determined the fitting residuals as the difference between the measured and the predicted DWI signal, where higher residuals correspond to lower quality of fitting (28). In general, we observed comparable fitting results between datasets A and B, while dataset C has slightly higher residuals across the brain. From the calculated fitting residuals in the whole-brain, we found that two subjects in Figure 6A (encircled in red) had high residuals across the three datasets, this could be due to motion artifact. On the other hand, in the WM ROIs, lower residuals were observed with dataset B compared to A and C, indicating possibility of lower signal in the acquisition with a b_*max*_ of 2000 s/mm^2^, rendering it less adequate for DKI fitting. In addition, although we employed the preprocessed (21) HCP dataset for the current study, subtle imaging artifacts (e.g., head movement, eddy-current, inhomogeneity of magnetic susceptibility, etc) could remain present in the underlying data that potentially affect the fitting quality. Furthermore, to check for any bias in our *in vivo* analysis, we also performed a pairwise correlation coefficient (r) test (see supplementary data Figures S2 - S4) for each subject between DKI parametric values from the three datasets. The findings suggested a high correlation (r~1) between all datasets, with consistent correlation between the b_*max*_ = 3000 s/mm^2^ datasets in both WM and GM. The DKI b_*max*_ dependencies observed in this study had been seen previously (14, 29), where approximation of signal intensity with DKI up to b-values of about 5000 s/mm^2^ had been examined. But the downside of deploying higher b-values is the reduced signal-to-noise ratio (SNR) for these data and lower reproducibility (14, 35), thus limiting the clinical feasibility of these acquisition approaches. In addition, the kurtosis method used here breaks down for b_*max*_ > 3000 s/mm^2^, where you would have to use higher orders of the cumulant expansion (i.e., up to b^3 instead of b^2) (14). Therefore, it’s been observed that better fitting accuracy is achieved up to b_*max*_ = 3000 s/mm^2^, while b_*max*_ > 3000 s/mm^2^ shows poor fitting quality (36). A major difference of this study, compared to previous work that evaluated DKI test-retest reliability, is the conclusion that adding another b-value shell between the protocol with b-value = 1000, 3000 s/mm^2^ does not provide much benefit. This also implies that a lower acquisition time can be achieved, increasing the potential of DKI as a diagnostic tool.

### 4.2 Verifying DKI reproducibility and fitting quality with phantom data

Finally, twelve tissue-mimicking phantoms grouped into four groups of three, representing distinct fiber orientations, (37) were incorporated in the study, which we used as ground truth data to further verify the DKI b_*max*_ dependency seen in the *in vivo* work. Similarly to the *in vivo* analysis of the whole-brain WM, selected WM bundles and GM, the reproducibility in the protocols with b_*max*_ = 3000 s/mm^2^ (datasets A and B) was higher in phantoms with fibers crossing at 30, 60 and 90 degrees. This indicates that b_*max*_ = 3000 s/mm^2^ could be needed to quantify varying microstructure complexity underlying different tissue types including WM fiber bundles of complex geometry. This is more evident in the MK, AK and RK maps. MK remains almost stable in terms of reproducibility across the different fiber crossings compared to the other maps, including DKI-derived DTI metrics (38). However, we also observed a lower variability in the 60° phantoms in all the datasets, which could be related to DKI’s ability to resolve fibers close to this configuration, but further analysis (outside the scope of the current study) is necessary to support this hypothesis.We have found the CoV for separately printed 3D phantoms with identical crossing angles to be between 2 and 8%, depending on the DKI parameter. Accordingly, the apparent dependence on crossing angle is likely a result in variabilities of the phantom manufacturing process. In addition, we assessed the goodness of fit in the ground truth data. These findings follow the same pattern as seen in our CoV results. The three shells dataset b-value = 1000, 2000, 3000 s/mm^2^ (dataset A) and two shells dataset b-value= 1000, 3000 s/mm^2^ (dataset B) maintained lower fitting residuals across the phantoms. On the other hand, higher residuals were seen in the other two shells dataset b-value = 1000, 2000 s/mm^2^ (dataset C). These findings indicate that DKI is robust for quantifying diffusion of water molecules reliably in heterogeneous microstructural environments while maintaining clinical feasibility.

## 5. LIMITATIONS

There are several limitations in the study. Our results may not be readily comparable to a more typical, clinical MRI setup. The current data are acquired using a modified 3T MRI system. This HCP scanner employs more powerful gradients to achieve the high spatial resolution (1.25mm isotropic) for the DWI data (21). Future work may investigate this potential beneficial effect of HCP’s custom hardware setup and inter-scanner variability. Moreover, the test-retest interval varies across subjects (average of 5 ± 3 months). However, we assume this effect on the kurtosis parameters reproducibility to be minimal as the brain’s structural configuration is not expected to change much within this time span. In contrast, phantoms were not moved between scans, hereby not changing partial volume effects and keeping CoVs low. Nevertheless, the phantom observations were in agreement with the *in vivo* results. Furthermore, the atlas-based ROIs and their ability to only capture the core part of the tracts may have influenced our WM ROI analysis. Finally, we employed the WLF algorithm since it was considered to give optimum results (14), therefore did not compare the effects of different fitting algorithms.

## 6. CONCLUSION

In conclusion, we have demonstrated that DKI reproducibility and quality of fitting depends on the maximum b-value used. Comparable reproducibility can be achieved with three (b-value = 1000, 2000, 3000 s/mm^2^) and the two (b-value = 1000, 3000 s/mm^2^) shell protocols, the latter having the advantage of shorter scan time. In contrast, the more common acquisition strategy (i.e., b-values =1000, 2000 s/mm^2^) is characterized by higher inter-scan variability. Although DKI has proven to be capable of characterizing non-Gaussian diffusion pattern in the brain (which is evident in ~90% of the WM voxels) (3), the inherent challenges of longer scan time compared to the traditional DTI might limit its potential use in clinical workflow. The test-retest reliability of DKI observed in this study with the b-value = 1000, 3000 s/mm^2^ dataset in both white and grey matter, and further verified on ground truth data, indicate that high reproducibility can still be achieved within a reasonable scan time, supporting DKI for clinical purposes. We propose that investigating the efficacy of the three protocols, especially the derived two shells (b-value = 1000, 3000 s/mm^2^) protocol in delineating subtle changes due to pathology in patient cohort, would be an appropriate future research direction.

## Acknowledgements

This work was supported by CIHR Foundation grant 201409, Canada Research Chairs Program 950-231964, NSERC Discovery Grant RGPIN-2015-06639, the Canada First Research Excellence Fund, Brain Canada, BrainsCAN and the Ontario Brain Institute Epilepsy Program (EpLink). The author Roy AM Haast is supported by BrainsCAN postdoctoral fellowships for this work.

## Supplementary Data

**Figure S1.**
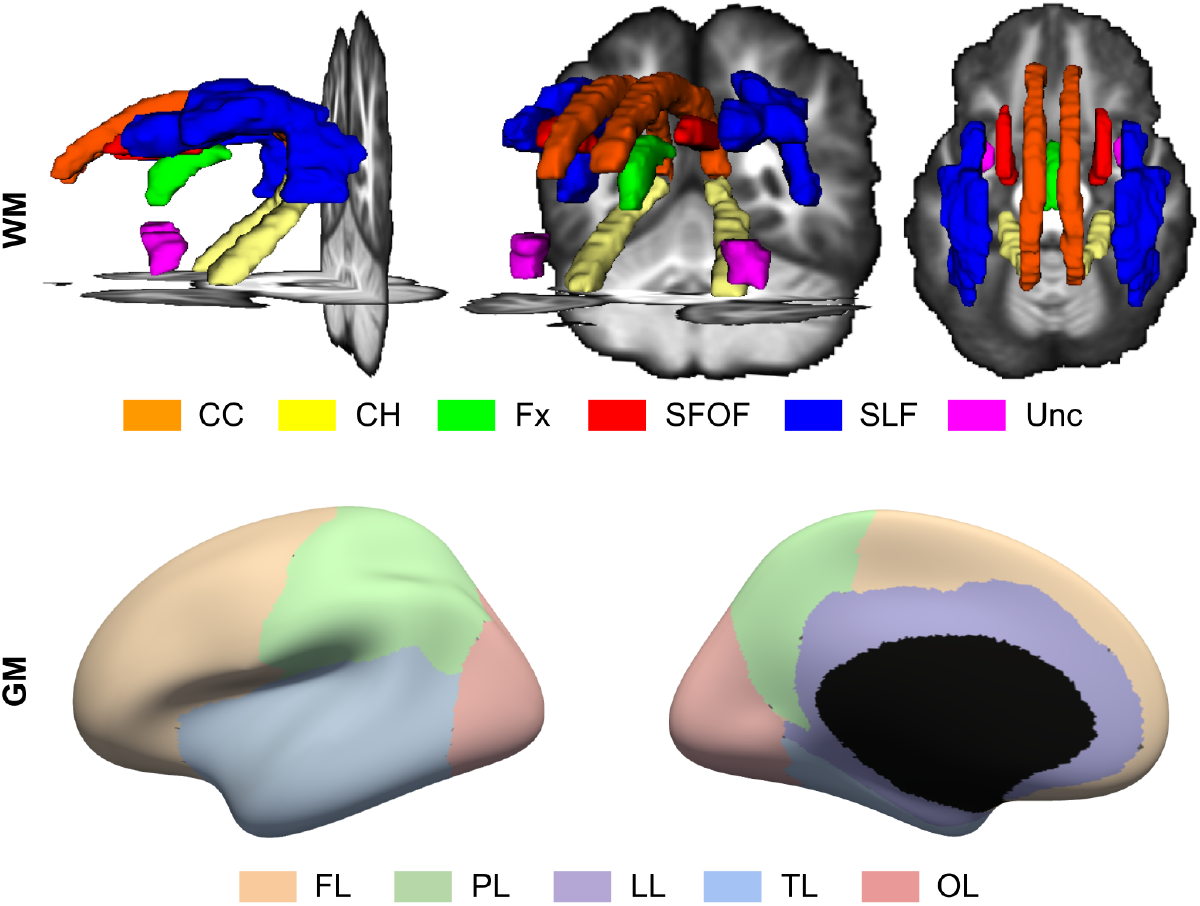
WM (top) and GM (bottom) ROIs used for analyses. Color-coded WM structures and GM regions indicate the different WM bundles (CC-Cingulum Cingulate, CH-Cingulum Hippocampus, Fx-Fornix, SFOF-Superior Fronto Occipital Fasciculus, SLF-Superior Longitudinal Fasciculus and Unc-Uncinate Fasciculus) as well as GM lobes (FL-Frontal Lobe, PL-Parietal Lobe, LL-Limbic Lobe, TL-Temporal Lobe and OL-Occipital Lobe).

**Figure S2.**
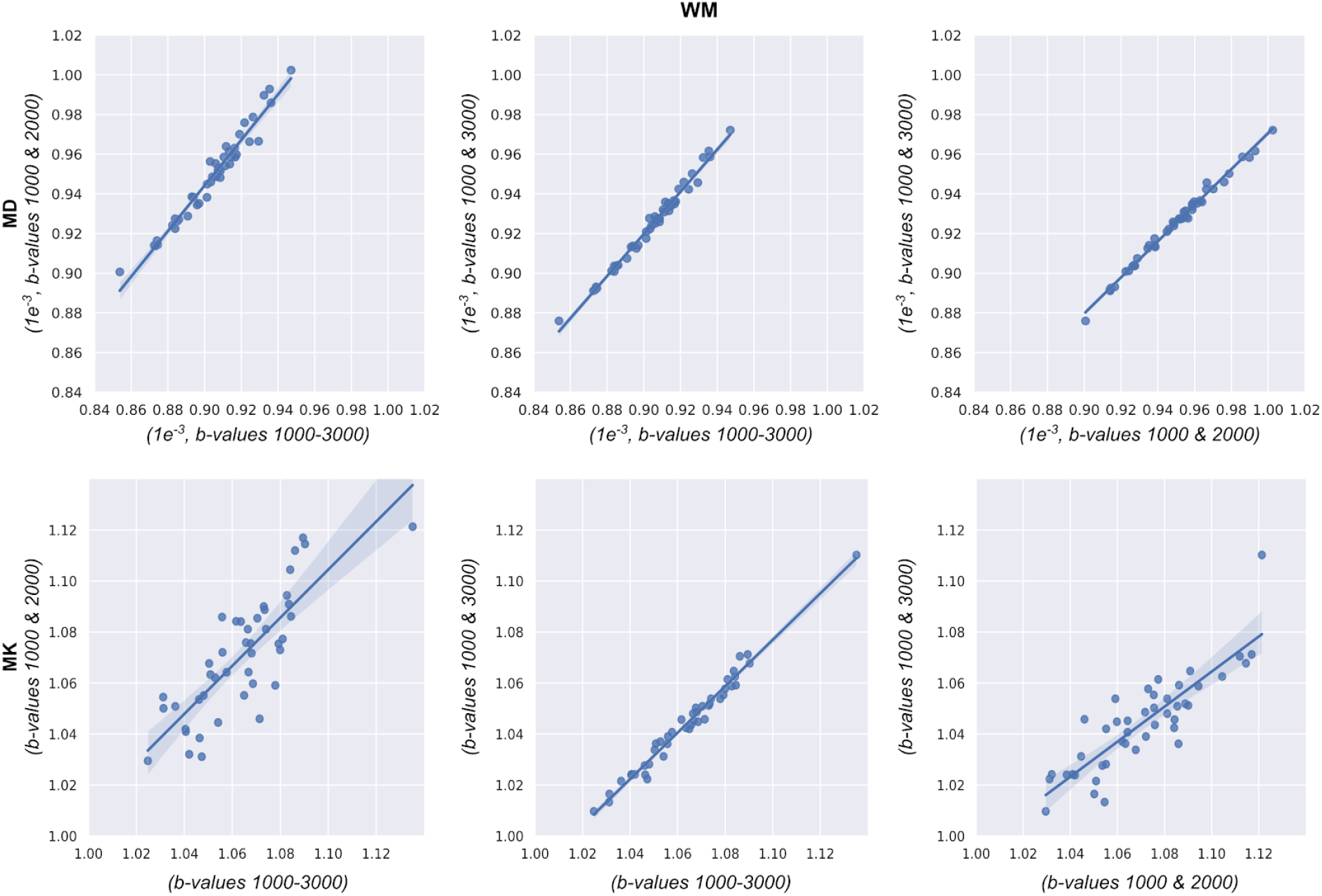
Pair-wise correlation coefficient (r) for each subject between DKI parametric values from the three datasets: A vs C, A vs B and C vs B (1^st^ row) for MD and A vs C, A vs B and C vs B (2^nd^ row) for MK. Shown are the individual subject voxel-wise values averaged across the whole-brain white-matter. There is a consistent high correlation between the three datasets (r ~1).

**Figure S3.**
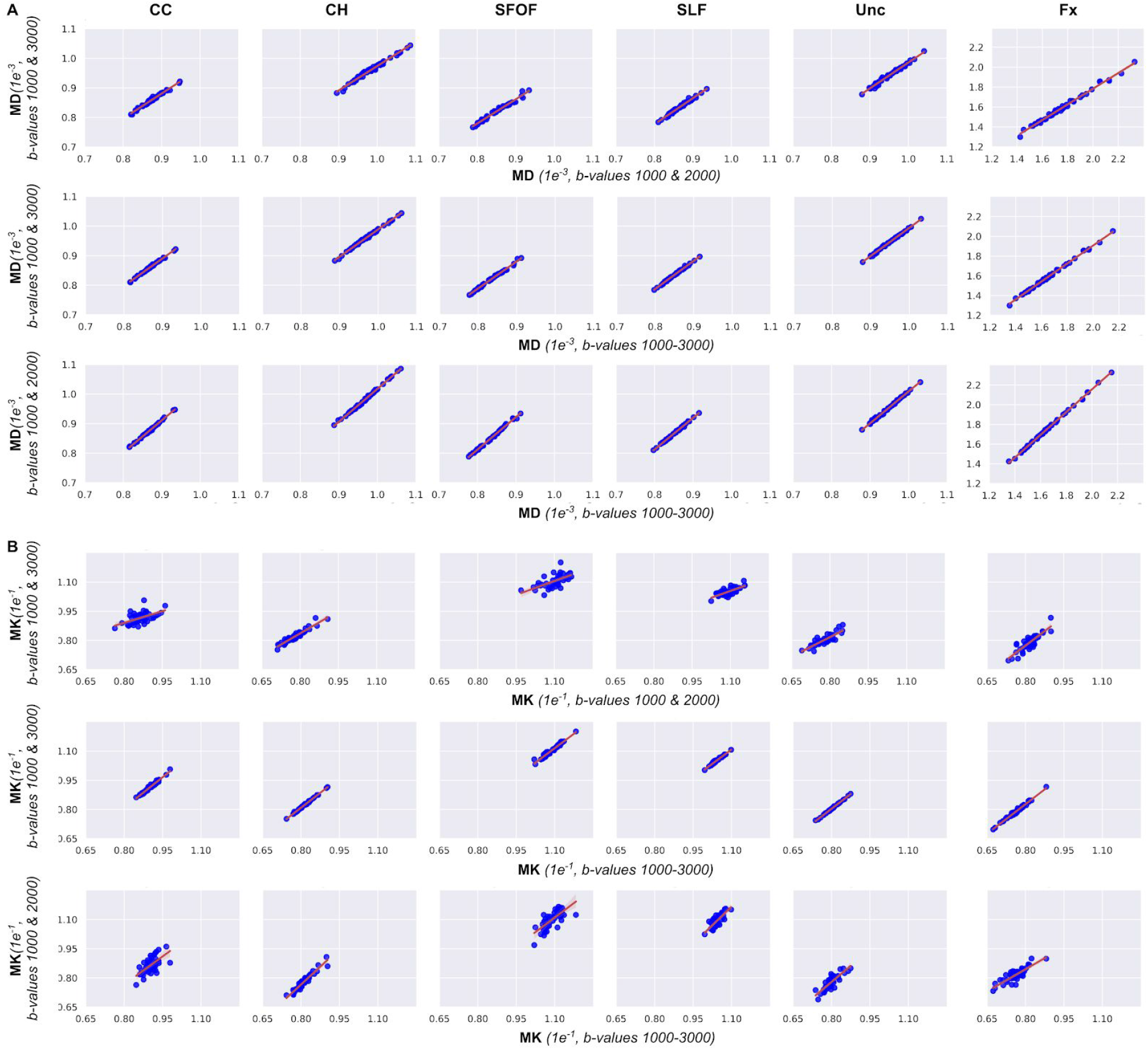
Pair-wise correlation coefficient (r) for each subject between DKI parametric values from the three datasets: C vs B (1^st^ row), A vs B (2^nd^ row) and A vs C (3^rd^ row) for MD and C vs B (4^th^ row), A vs B (5^th^ row) and A vs C (6^th^ row) for MK. Shown are the individual subject voxel-wise values averaged across the white-matter ROIs. There is a consistent high correlation between the three datasets (r ~1).

**Figure S4.**
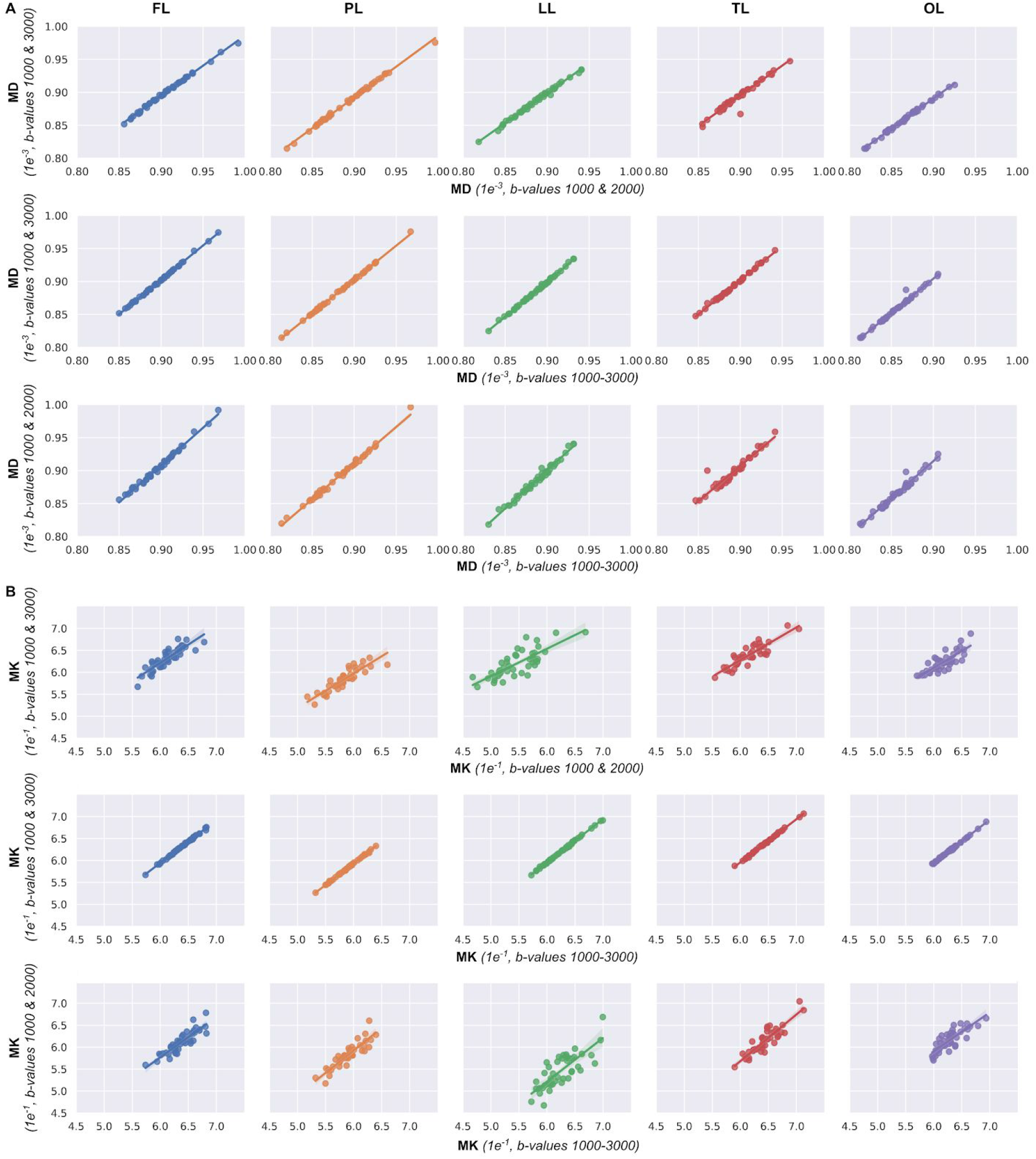
Pair-wise correlation coefficient (r) between DKI parametric values from the three datasets: C vs B (1^st^ row), A vs B (2^nd^ row) and A vs C (3^rd^ row) for MD and C vs B (4^th^ row), A vs B (5^th^ row) and A vs C (6^th^ row) for MK. Shown are the individual subject vertex-wise values averaged across each lobe of the left hemisphere (similar results observed in the right hemisphere). For all the lobes there is a consistent high correlation (r~1) between the three datasets.

**Table S1.** Mean DKI values for each of the phantoms representing different fiber orientations (in degrees), shown for MK and MD only. The Pearson correlation was high between the three datasets (A, B and C, r~1)., Except for MD from dataset C, where we observed low correlation with the 60° phantom.

| Map | Fiber orientation | B-values |  |  |
| --- | --- | --- | --- | --- |
|  |  | 1000-3000 | 1000 & 3000 | 1000 & 2000 |
| MD | 0 | 0.136 | 0.135 | 0.137 |
| MD | 30 | 0.056 | 0.053 | 0.063 |
| MD | 60 | 0.025 | 0.028 | 0.002 |
| MD | 90 | 0.029 | 0.03 | 0.012 |
| MK | 0 | 0.136 | 0.138 | 0.132 |
| MK | 30 | 0.136 | 0.136 | 0.135 |
| MK | 60 | 0.134 | 0.134 | 0.127 |
| MK | 90 | 0.137 | 0.138 | 0.127 |

## Notes

### Competing Interest Statement

The authors have declared no competing interest.

